# Advancing insect vector biology research: a community survey for future directions, research applications and infrastructure requirements

**DOI:** 10.1101/042242

**Authors:** Alain Kohl, Emilie Pondeville, Esther Schnettler, Andrea Crisanti, Clelia Supparo, George K. Christophides, Paul J. Kersey, Gareth L. Maslen, Willem Takken, Constantianus J. M. Koenraadt, Clelia F. Oliva, Núria Busquets, F Xavier Abad, Anna-Bella Failloux, Elena A. Levashina, Anthony J. Wilson, Eva Veronesi, Maëlle Pichard, Sarah Arnaud Marsh, Frédéric Simard, Kenneth D. Vernick

**Affiliations:** MRC-University of Glasgow Centre for Virus Research, Glasgow G61 1QH, Scotland, UK; Department of Life Sciences, Imperial College London, London SW7 2AZ; The European Molecular Biology Laboratory - The European Bioinformatics Institute, Wellcome Trust Genome Campus, Hinxton, Cambridge CB10 1SD, UK; Laboratory of Entomology, Wageningen University and Research Centre, P.O. Box 16, 6700 AA Wageningen, The Netherlands; Polo d’Innovazione di Genomica, Genetica e Biologia, P.zza Gambuli, Edificio D, 3° Piano, 06132 Perugia, Italy; Centre de Recerca en Sanitat Animal (CReSA)—Institut de Recerca i Tecnologia Agroalimentàries (IRTA), Campus UAB, 08193 Bellaterra, Barcelona, Spain; Arboviruses and Insct Vectors Unit, Department of Virology, Institut Pasteur, 25-28 rue du Docteur Roux, 75724 Paris cedex 15, France; Department of Vector Biology, Max-Planck-Institut für Infektionsbiologie, Campus Charité Mitte, Charitéplatz 1, 10117 Berlin, Germany; Integrative Entomology Group, Vector-borne Viral Diseases Programme, The Pirbright Institute, Ash Road, Pirbright, Woking, Surrey GU24 0NF, UK; Swiss National Centre for Vector Entomology, Institute of Parasitology, University of Zürich, 8057 Zürich, Switzerland; Department of Parasites and Insect Vectors, Institut Pasteur, Unit of Insect Vector Genetics and Genomics, 28 rue du Docteur Roux, 75015 Paris cedex 15, France; MIVEGEC “Maladies Infectieuses et Vecteurs: Ecologie, Génétique, Evolution et Contrôle”, UMR IRD224-CNRS5290-Université de Montpellier, 911 Avenue Agropolis, 34394 Montpellier, France; CNRS Unit of Hosts, Vectors and Pathogens, Paris, France (URA3012), 28 rue du Docteur Roux, 75015 Paris cedex 15, France

## Abstract

**Background:** Vector-borne pathogens impact public health and economies worldwide. It has long been recognized that research on arthropod vectors such as mosquitoes, ticks, sandflies and midges which transmit parasites and arboviruses to humans and economically important animals is crucial for development of new control measures that target transmission by the vector. While insecticides are an important part of this arsenal, appearance of resistance mechanisms is an increasing issue. Novel tools for genetic manipulation of vectors, use of *Wolbachia* endosymbiotic bacteria and other biological control mechanisms to prevent pathogen transmission have led to promising new intervention strategies. This has increased interest in vector biology and genetics as well as vector-pathogen interactions. Vector research is therefore at a crucial juncture, and strategic decisions on future research directions and research infrastructures will benefit from community input.

**Methodology/Principal Findings:** A survey initiated by the European Horizon2020 INFRAVEC-2 consortium set out to canvass priorities in the vector biology research community and to determine key issues that should be addressed for researchers to efficiently study vectors, vector-pathogen interactions, as well as access the structures and services that allow such work to be carried out.

**Conclusions/Significance:** We summarize the key findings of the survey which in particular reflect priorities in European countries, and which will be of use to stakeholders that include researchers, government, and research organizations.

**Author Summary:** Research on arthropod vectors that transmit so-called arboviruses or parasites, such as mosquitoes, ticks, sandflies and midges is important for the development of control measures that target transmission of these pathogens. Important developments in this research area, for example vector genome sequencing, genome manipulation and use of transmission-blocking endosymbionts such as *Wolbachia* have increased interest in vector biology. As such, strategic decisions on research directions as well as research infrastructures will benefit from community input. A survey initiated by the European Horizon2020 INFRAVEC-2 consortium set out to investigate priorities in the vector biology research community as well as key issues that impact on research, and access to the structures and services that allow such studies to be carried out. Here we summarize the key findings of this survey, which in particular reflect priorities in European countries. The survey data will be of use to decision makers such as governments and research organizations, but also researchers and others in the field.

## Introduction

Vector-borne diseases such as those transmitted by mosquitoes have a major impact on human and animal health. Among the many examples, malaria (caused by *Plasmodium* parasites) and dengue (caused by four serotypes of dengue virus, *Flaviviridae*) stand out as major diseases that affect populations worldwide, but new threats such chikungunya virus (*Togaviridae*) and more recently Zika virus (*Flaviviridae*) have emerged [1-4]. Both known and emerging pathogens put huge pressure on communities and public health systems. Vaccine development against key threats to human health such as dengue virus and *Plasmodium* parasites may offer tools against transmission and disease, and progress is encouraging [5-8]. However, issues such as pathogen strain variation and vaccine or drug production/distribution costs will remain as challenges [9], and even with vaccines vector control will be a crucial part of a multivalent arsenal. Although drugs against malaria parasites are on the market, availability, administration and resistance are problematic [10-12]. Drugs targeting dengue virus are in the development stages [13-16]. In the case of chikungunya virus vaccine candidates and drugs are now in development [17]. Only veterinary vaccines are currently in use for the animal pathogen Rift Valley fever virus (*Bunyaviridae*) and efforts to produce human vaccines are urgently needed [18, 19].

Many ongoing efforts to control vector-borne diseases rely on control measures that target mosquitoes including control of larval breeding sites, use of insecticides, use of bed nets (often used in combination with insecticides) (see for example, [20-25]). These efforts have been successful when implemented consistently, although issues such as insecticide resistance, changes in vector behavior, and difficulties with breeding site control (see for example [26-35]) require that research in vector biology and control is continuously developed and strengthened. Technological developments over the last decade are transforming modern vector research. These include: vector genome sequences, high-throughput genomics, transcriptomics, and population genetics with results in public databases [36], improved methods for genetic manipulation of arthropods (that have led to field trials) [37-43], studies on the influence of the mosquito midgut microbiome on pathogen transmission [44-46], studies on the impact of the insect-specific viruses on arbovirus transmission [47, 48], and the use of *Wolbachia* endosymbiotic bacteria that prevent pathogen transmission [37, 49-52]. Nonetheless the opportunity to access and make best use of ongoing research can be difficult, given the specialized knowledge, costs and infrastructures required.

The European Union (EU) has identified access to specialized Research Infrastructures (RIs) as a key to producing high quality science. RIs are defined as “Tools for science….RIs offer **unique research services** to users from different countries, attract young people to science, and help to shape scientific communities…‥ RIs may be ‘**single-sited**’ (a single resource at a single location), ‘**distributed**’ (a network of distributed resources), or ‘**virtual**’ (the service is provided electronically)” [53]. Such RIs can be research facilities, resources and related services. Within the Framework Programmes (FP) of the EU, Research Infrastructure projects support the improvement of key high-level facilities for research, and allow access to the facilities by researchers in Europe and eligible member states. A wide range of research disciplines have been targeted by RI projects, including physics, information science, earth science and medicine. One such Research Infrastructure project under EU FP7 was Infravec, which focused on developing and providing research resources for insect vector biology from 2009-2014. Infravec, which was constituted as an EC Starting Community under FP7, obtained the opportunity to renew the project as an Advanced Community (AC) called INFRAVEC-2 under the Horizon 2020 framework. Research Infrastructure projects are not research networks, but rather are tasked to identify the key unique and rare research infrastructures necessary for a research community, and organize them so that researchers at institutes lacking the Research Infrastructures can access the facilities in order to expand the scope of their research. Thus, Research Infrastructures are exceptional facilities that permit experiments that could not routinely be done without this structure. Use of Research Infrastructure facilities by external researchers is provided as so-called “Transnational Access” (TNA), with access costs reimbursed by the Research Infrastructure project, thus provided at no cost to the end-user.

Conditions have changed since the inception of the FP7 Infravec project, including the emergence and transmission of arboviruses in Europe and elsewhere, as well as widening the project scope to include vector-borne diseases of economically important animals and the most recently developed innovative technologies. Collecting information about the current and perceived future infrastructure needs of the vector biology research community and other stakeholders is an important step to ensure that the services offered via Transnational Access reflect actual needs of the advanced community. Here we present the findings of a survey of scientists and associated stakeholders in the field of vector biology or fields that are linked to vector biology such as pathogen studies, which will help to define priorities and requirements within INFRAVEC-2 but should also be of interest to governments, research organizations and researchers in the field. Participation numbers suggest that in particular European research priorities are reflected in the results, but the data can inform stakeholders worldwide.

## Materials and Methods

### Survey structure

A questionnaire (S1 Table) was sent to organizational email lists (European Society for Vector Ecology; the journal Pathogens and Global Health; National Center of Expertise in Vectors (CNEV, France); CIRM-Italian Malaria Network; FP7 Infravec mail list; International Meeting on Arboviruses and their Vectors mail list; BioInsectes; EU/DEVCO MEDILABSECURE network), as well as to other lists owned by the authors. The questionnaire was sent as a URL link to the online form along with an explanatory note to scientists in the vector biology field and associated stakeholders. The questionnaire request was spontaneously retransmitted by an unknown number of recipients to organizational and other lists.

Briefly, the cover note explained the aims of the INFRAVEC-2 community, followed by a series of questions. The key areas covered by the survey are as follows: 1) vectors and vector-borne pathogens studied by survey participants, 2) research area (with several responses allowed), 3) infrastructures available at the respondents home institution including those for vector and animal research, 4) ease of access to vector research facilities outside the survey participants’ home institution, 5) infrastructures that participants would use, offered by the facilities at no cost to user, 6) identification of research priorities over the next 5-10 years, and 7) additional feedback. The survey was carried out from October to November 2015. Respondents were given the opportunity to provide their name and institution, although this was not required for completion of the questionnaire. However, all respondents (n=211) identified themselves, indicating that repeat voting or vote stuffing is not a concern for interpretation of the results. All results shown here are anonymized, and no survey participant details published.

## Results and Discussion

In total 211 responses were obtained (see S2 Table). Approximately 88% of respondents were from countries across Europe, with France, and then the UK providing the highest numbers of responses. This suggests that the results reflect a good overview of current priorities in European vector biology and vector research areas. Below we summarize and analyze the data obtained in the survey.

## Research areas: arthropods and pathogens relevant to survey participants

Our goal was to obtain an overview of the research areas and work of survey participants, which are thus likely to guide their future research needs (S1 Table, Survey Questionnaire). First, respondents indicated vectors relevant to their research as major or minor area of interest (Table 1). *Aedes* species mosquitoes were the top field, followed by *Anopheles* and *Culex* species. The strong interest in aedine species may reflect the emergence of arboviruses such as chikungunya transmitted by *Ae. aegypti* and *Ae. albopictus* as well as the expansion of the latter species in Europe (and other areas) and acting as arbovirus vector [54-60]. Despite their importance in the European context as major vectors of pathogens, comparatively little research is carried out on ticks and *Culicoides* midges. This suggests that the European vector biology community presently lacks sufficient opportunities and resources for research on these vectors. Among the category “Other”, comments by participants indicated phlebotomines/sand flies as a key area, with tsetse flies, fleas, triatomines and tabanids/horse flies also mentioned.

**Table 1.** Research areas and interests of the survey participants. Numbers of responses are indicated as Major or Minor depending on vector listed, or in the category “Other” which incorporates other vectors not specifically listed (selection of responses shown).

| Arthropod | Major | Minor |
| --- | --- | --- |
| *Aedes* spec. | 102 | 38 |
| *Culex* spec. | 66 | 46 |
| *Anopheles* spec. | 79 | 42 |
| *Culicoides* spec. | 24 | 32 |
| Ticks | 55 | 44 |
| Other | 42 | 49 |
| Vectors mentioned under “Other” (selection of most mentioned): phlebotomines/sandflies (33), fleas (13), tsetse flies (7), triatomines (4), tabanids/horse flies (6) |  |  |

We also quantified the major and minor interests of survey participants (Table 2). There was a notably strong indication of research interests in arboviruses, mainly affecting humans but also livestock pathogens as well. These research interests and activities are likely due to the emergence and importance of arboviruses such as chikungunya, Zika, Schmallenberg and bluetongue [2, 55, 61-63]. Given the historically important role of malaria research also in Europe, the overall importance in the vector field is not surprising. Of note was the impact of tick-borne pathogens in the category “Other” and this is worth mentioning especially with the impact of Lyme disease across Europe and North America [64] and surge in interest in Crimean-Congo hemorrhagic fever virus [65, 66].

**Table 2.**
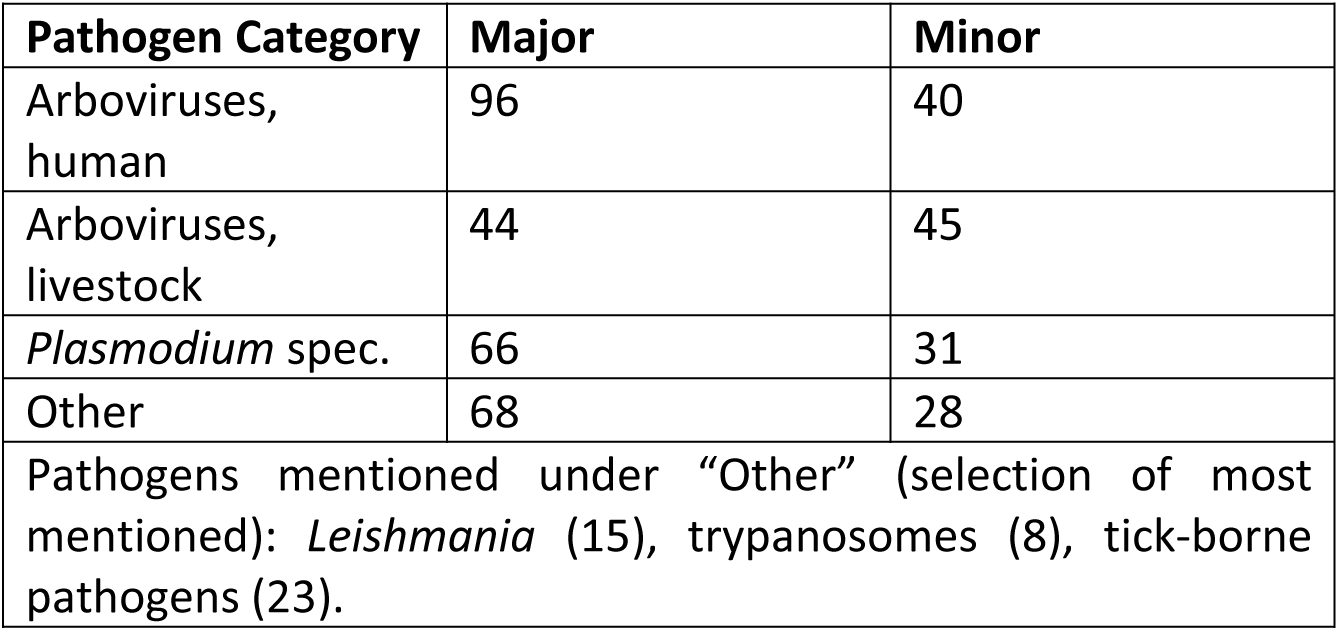
Pathogens relevant to the survey participants. Numbers of responses are indicated as Major or Minor depending on pathogen category listed, or in the category “Other” which incorporates other pathogens not specifically listed (selection of responses shown).

To describe their activities in more detail, we collected further data on the research areas of interest to the survey participants (Table 3). In general vector biology describes the research of over half of the participants, however this is a very broad term. Vector ecology, behavior and control were also commonly reported. Of note, genetic modification and vector immunity remain relatively small fields despite important advances in these areas. Interest may increase with better tools and access to new resources such as strains and facilities. The survey data showed that studies of pathogens either directly or within the context of host-pathogen or vector-pathogen interactions are a key area of research. This needs to be emphasized as it integrates disciplines such as virology, parasitology, cell biology, microbiology and genetics into the vector field. Similarly, surveillance, diagnostics and epidemiology were important areas and this (alongside vector control, behavior and ecology) was an indication of the applied character of many activities in the field of vector-borne diseases.

**Table 3.**
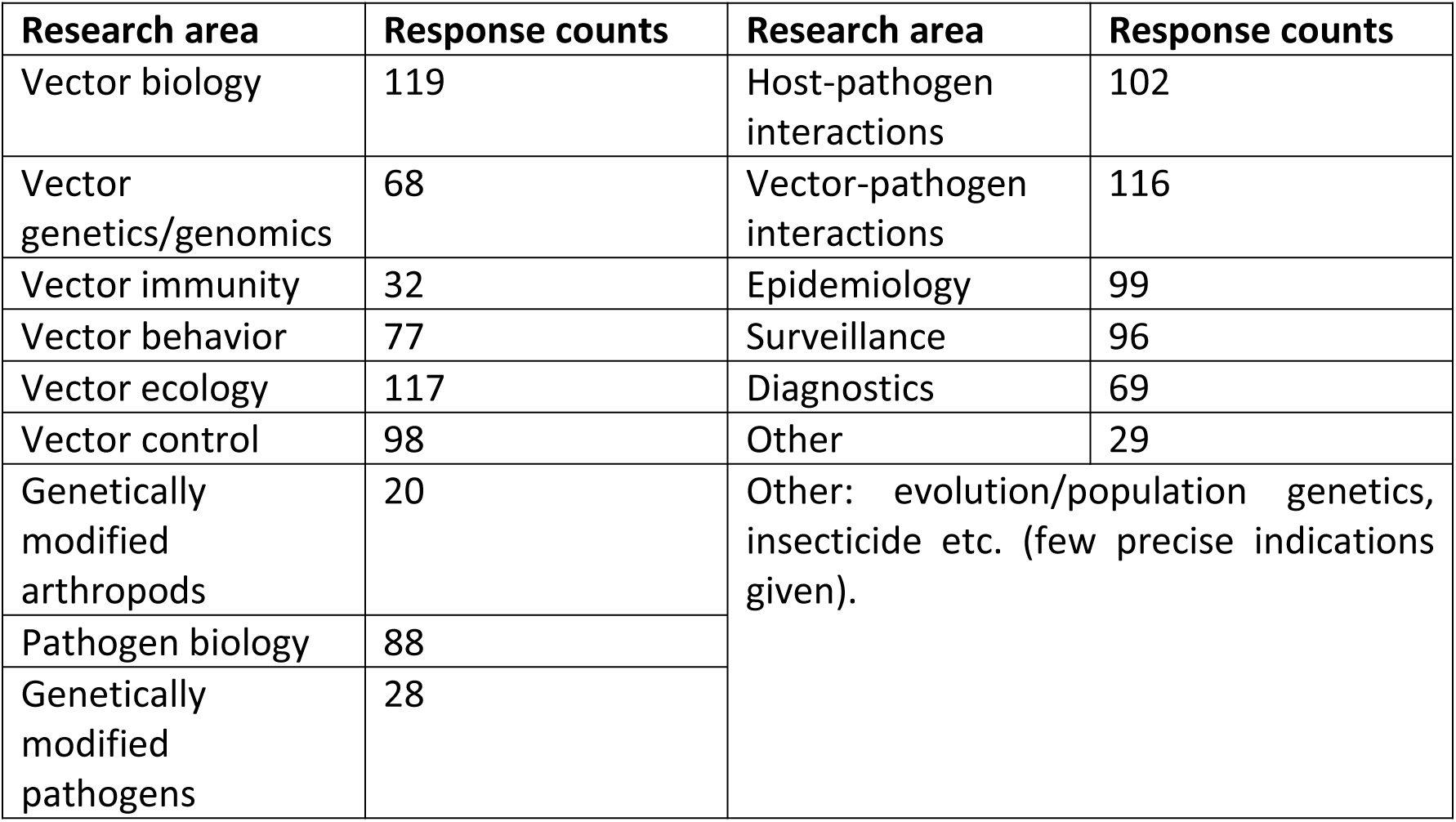
Details of research areas relevant to survey participants. Numbers of responses are shown by research area, or in the category “Other” which incorporates fields not specifically listed (selection of responses shown).

| Research area | Response counts | Research area | Response counts |
| --- | --- | --- | --- |
| Vector biology | 119 | Host-pathogen interactions | 102 |
| Vector genetics/genomics | 68 | Vector-pathogen interactions | 116 |
| Vector immunity | 32 | Epidemiology | 99 |
| Vector behavior | 77 | Surveillance | 96 |
| Vector ecology | 117 | Diagnostics | 69 |
| Vector control | 98 | Other | 29 |
| Genetically modified arthropods | 20 | Other: evolution/population genetics, insecticide etc. (few precise indications given). |  |
| Pathogen biology | 88 |  |  |
| Genetically modified pathogens | 28 |  |  |

## Assessment of currently available facilities

Knowledge of availability and/or ease of access to research infrastructures is a key factor in future planning of research activities. Survey participants were therefore asked to indicate their current organization’s current capabilities. As shown in Table 4, survey participants indicated a certain level of capacity to provide vectors but also material across the community. Moreover facilities for biosafety level (BSL) 2 and 3 experiments with vectors, animals and pathogens are available in several places. The concept of Research Infrastructure can be extended to reagent provision and has been successfully established by FP7 Infravec and the European Virus Archive (http://www.european-virus-archive.com). This indicates an existing infrastructure base that can be developed and made available for research on vectors and pathogens on a wider basis (for example those who do not have immediate access to BSL 3 level insectaries but would require experiments to be carried out in such facilities) through communities such as INFRAVEC-2.

**Table 4.** Research infrastructures and resources available to survey participants. Various types of structures relevant to vector and pathogen research are indicated.

| Available facilities and resources | Response counts |
| --- | --- |
| Furnish vectors to external users | 74 |
| Furnish BSL2/BSL3 infected vectors/extracts to external users | 32 |
| BSL2 containment: arthropod infections | 91 |
| BSL3 containment: arthropod infections | 60 |
| Pathogen work in cell culture | 128 |
| BSL2 or BSL3 containment: small animal work | 83 |
| BSL2 or BSL3 containment: large animal work | 27 |

## Assessment of infrastructure and service requirements

When survey participants were asked to indicate how many had requested access to insectaries at BSL2 or 3 in other institutions, in total 62 positive responses were received. However out of these, 18 responses indicated that access could not be granted in a timely manner. This suggests that inability to consistently access secure insectary facilities comprises a systematic weakness that impedes research on vector-pathogen interactions and may also explain the weaker interest in vector immunity studies, for example. The relevant secure insectary facilities exist in Europe (Table 4), and thus a mutualized network of insectaries at BSL2 and 3 could resolve access limitations and promote elevated levels of vector research under BSL2 and 3 conditions.

Access needs, or provision of infected vectors or extracts from infected vectors were assessed and participants were asked to indicate which pathogens or facilities/services would be of interest in the context of INRFAVEC-2 where these are free of cost (or the requirement for collaboration) for the end user (Table 5). Although the questions below were originally aimed at potential European users all answers were taken into account. Survey data show that in particular services and structures for arbovirus research would likely generate strong demand. Again this may be due to the surge in research in this field described above. Similarly, BSL2 and 3 studies on infected vectors and insecticides as well as behavior scored highly. Regarding technologies novel for the field, functional siRNA screens and imaging of vectors did not score particularly high but this demand may increase in the future, particularly if facilities were available for access.

**Table 5.**
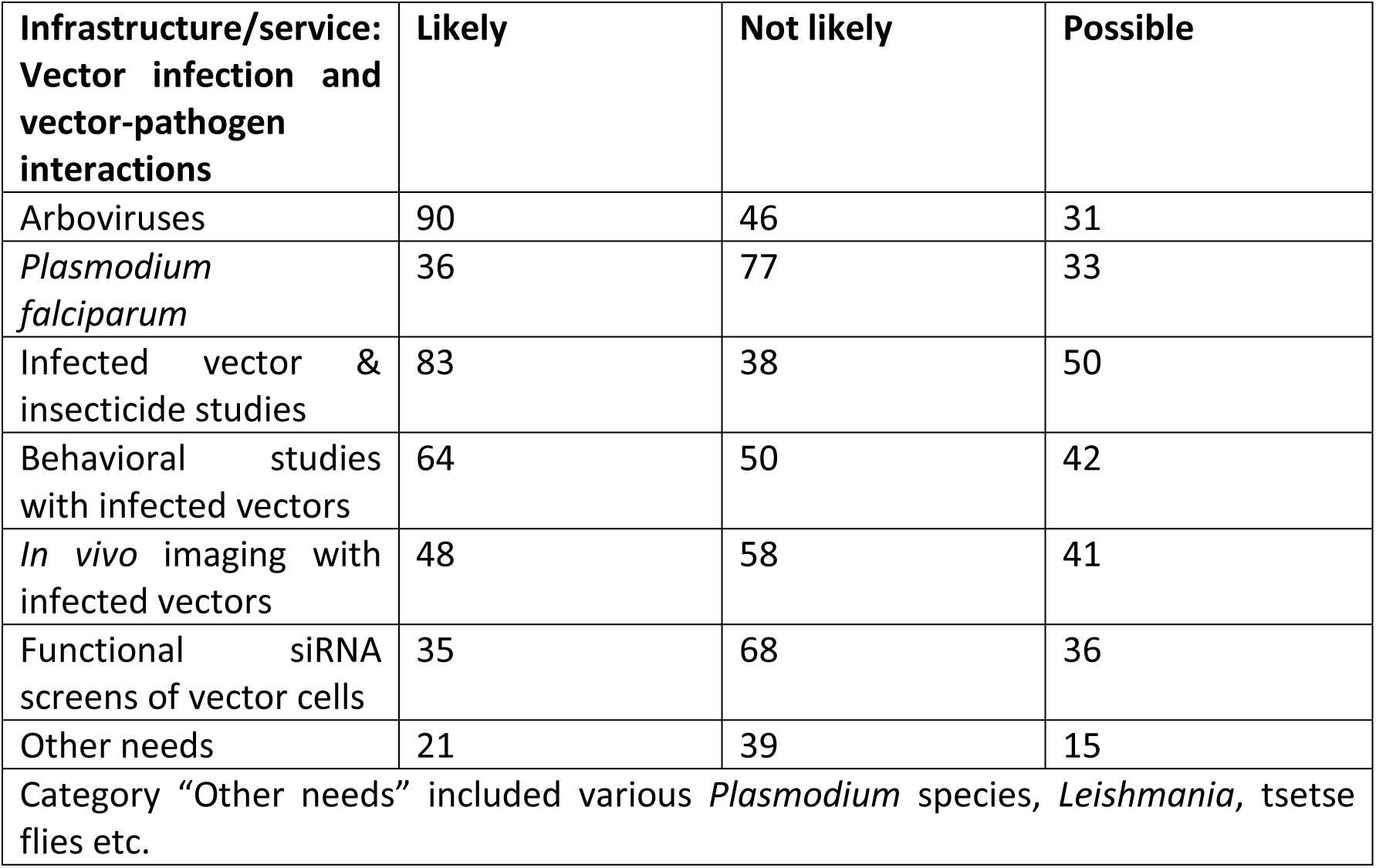
Infrastructure services (vector infection and vector-pathogen interactions) for the vector research community. Survey participants responded whether the services listed here (vector infection and vector-pathogen interactions) to study vector infections and vector-pathogen interactions, would be of use if offered free of cost. Response counts are grouped into Likely, Not likely or Possible use of the infrastructure/service.

| Infrastructure/service:<br>Vector infection and<br>vector-pathogen<br>interactions | Likely | Not likely | Possible |
| --- | --- | --- | --- |
| Arboviruses | 90 | 46 | 31 |
| *Plasmodium falciparum* | 36 | 77 | 33 |
| Infected vector &<br>insecticide studies | 83 | 38 | 50 |
| Behavioral studies<br>with infected vectors | 64 | 50 | 42 |
| *In vivo* imaging with<br>infected vectors | 48 | 58 | 41 |
| Functional siRNA<br>screens of vector cells | 35 | 68 | 36 |
| Other needs | 21 | 39 | 15 |
| Category "Other needs" included various *Plasmodium* species, *Leishmania*, tsetse flies etc. |  |  |  |

Vector genetics and genomics (see www.vectorbase.org, [36]) but also studies of vector microbiomes (given their influence on mosquito infection with arboviruses and parasites [44-46, 67]) are expanding fields. These research areas have strongly benefited from high-throughput sequencing techniques and bioinformatics. Survey participants were enthusiastic about developing insect vector-oriented infrastructures, services and expertise in high throughput genomics and bioinformatics, especially transcriptional profiling and genome and population analysis (Table 6).

**Table 6.** Infrastructure services (vector genomics and bioinformatics) for the vector research community. Survey participants responded whether the services listed here (vector genomics and bioinformatics), would be of use if offered free of cost. Response counts are grouped into Likely, Not likely or Possible use of the infrastructure/service.

| Infrastructure/service:<br>Vector genomics and<br>bioinformatics | Likely | Not likely | Possible |
| --- | --- | --- | --- |
| Transcriptional<br>profiling | 75 | 42 | 53 |
| Genome or population<br>analysis | 72 | 43 | 54 |
| Bacterial microbiome<br>profiling | 45 | 63 | 41 |
| Population or focused<br>SNP genotyping | 39 | 63 | 48 |
| Other needs | 10 | 41 | 8 |
| Category "Other needs" included proteomics, metabolomics. |  |  |  |

The era of genomics has brought about much needed information on vector genomes (see for example [68-70]). Genetic manipulation of genomes in basic biological studies of gene/sequence structure and function, and applications based on genome manipulation (see for example [71-73]) are useful tools to maximize the value of this information, and for example CRISPR/Cas9-mediated genome manipulation is an important technical advance also for the vector field [74, 75]. We therefore asked survey participants about their interest in applying genome editing technologies within their work. As shown in Table 7, there was particularly strong interest in genetic manipulation of aedine mosquitoes. *Culicoides* midges seemed at present a less popular subject, probably at least in part because the community is small as mentioned above, as well as that the technologies have not yet been applied to this system or general issues with establishing colonies of important midge vector species. Among the category “Other”, ticks stood out.

**Table 7.** Infrastructure services (vector genome editing) for the vector research community. Survey participants responded whether vector genome editing would be of use if offered free of cost. Response counts are grouped into Likely, Not likely or Possible use of the infrastructure/service.

| Infrastructure/service:<br>Vector genome editing | Likely | Not likely | Possible |
| --- | --- | --- | --- |
| *Anopheles* spec. | 35 | 71 | 44 |
| *Aedes* spec. | 60 | 60 | 33 |
| *Culicoides* spec. | 20 | 85 | 17 |
| Other | 27 | 53 | 7 |
| "Other" included ticks (17), phlebotomines (5), *Culex* spec. and tsetse flies (both 4). |  |  |  |

Studies on vectors (infected, uninfected or genetically modified) often include components that analyze behavior and ecology. A further section of this survey therefore focused on a number of specific potential requirements in this area. As indicated in Table 8, the interest to work in field sites in endemic countries if access could be provided, as well as standardized behavioral assays and bioassays for vectors generated strong positive responses. This suggested a need for these in the vector research community. Positive responses for large cage studies (controlled indoors or semi-controlled outdoors) were also strong considering that such applications are very specialized, and the facilities are rare. However, this illustrates the potential contribution of a Research Infrastructure project, because community mutualization of rare infrastructures can allow access to state of the art facilities for researchers with occasional needs. In the future, the possibility to access such facilities may become stronger as more genetically modified vectors will be assessed in pre-release assays. Few positive responses for electrophysiology experiments were obtained, suggesting that there is no major need for additional facilities beyond what is already in place.

**Table 8.** Infrastructure services (vector ecology and behavior) for the vector research community. Survey participants responded whether specific services or infrastructures to study vector ecology and behavior, would be of use if offered free of cost. Response counts are grouped into Likely, Not likely or Possible use of the infrastructure/service.

| **Infrastructure/service:**<br>**Vector ecology and**<br>**behavior** | **Likely** | **Not likely** | **Possible** |
| --- | --- | --- | --- |
| Facilitated work at endemic country field sites | 96 | 39 | 43 |
| Electrophysiology | 14 | 99 | 23 |
| Standardized vector behavioral assays & bioassays | 65 | 52 | 52 |
| Large cage studies (controlled large indoor insectary) | 64 | 61 | 46 |
| Large cage studies (semi-controlled outdoor large cages) | 46 | 77 | 40 |
| Other needs | 7 | 44 | 3 |
| Very few responses to “Other needs” given, for example cage trials in Europe. |  |  |  |

Survey participants were also asked about their requirements for more specific vector-related data and research resources such as reference genomes, specific cell lines and mosquito strains (Table 9). Results indicated that in particular, a bank of standard vector colonies would be of interest to the community. Easily accessible quality-controlled vector colonies available from a European repository could be an important influence promoting comparability and reproducibility of experimental infection and other results across laboratories. Similarly vector systematics and collections generated high interest. However, the practices of systematics may be at a juncture, because the technological capacity will soon be available to whole-genome sequence large numbers of unidentified individuals of a putative vector clade, and cluster them bioinformatically to determine phylogenetic relatedness. These results will need to be compared to existing collections, including voucher specimens. Perhaps surprisingly, new reference and cloned vector cell lines did not score highly but these may be of interest to smaller research areas such as virologists who carry out particular types of studies. Cell lines may be of less interest in malaria vector research where the biology is not consistent with simple cell models of *Plasmodium*-mosquito interaction. Despite high interest in *Wolbachia* to block pathogen transmission [51], generation of novel trans-infected vector strains was also not a priority. Finally, a small number of responses under “Other needs” mentioned the importance of training.

**Table 9.** Infrastructure services (vector biology resources) for the vector research community. Survey participants responded whether specific resources for vector biology, would be of use if offered free of cost. Response counts are grouped into Likely, Not likely or Possible use of the infrastructure/service.

| **Infrastructure/service:**<br>**Vector biology**<br>**resources** | **Likely** | **Not likely** | **Possible** |
| --- | --- | --- | --- |
| Bank of standard vector reference strains (genome & RNA sequenced) | 85 | 34 | 54 |
| Colonization of novel vector strains & species | 76 | 45 | 53 |
| Production of new reference vector cell lines (genome & RNA sequenced) | 38 | 74 | 38 |
| Production of cloned vector cell lines | 39 | 75 | 37 |
| Production of microbiome-free mosquitoes | 28 | 82 | 40 |
| *Wolbachia*-transinfected vector strains | 23 | 76 | 52 |
| Vector systematics and collections | 62 | 52 | 55 |
| Other needs | 5 | 44 | 3 |
| Very few responses to "Other needs" given, mainly mentioning training needs. |  |  |  |

Our survey specifically addressed training needs, community networking and communication. As shown in Table 10, all suggestions - training in vector BSL2 and 3 methods, training in bioinformatics and genomics and scientific communication by conferencing - were positively received by the survey participants. Clearly these are areas of need that should be developed as a real requirement within the vector research field.

**Table 10.** Infrastructure services (training and networking activities) for the vector research community. Survey participants responded whether specific services or infrastructures in the areas of training and networking activities, would be of use if offered free of cost. Response counts are grouped into Likely, Not likely or Possible use of the infrastructure/service.

| **Infrastructure/service:**<br>**Training and**<br>**networking activities** | **Likely** | **Not likely** | **Possible** |
| --- | --- | --- | --- |
| Training in BSL2 and 3 vector infection and study techniques | 96 | 30 | 51 |
| Training in bioinformatics and genomic analysis | 107 | 25 | 53 |
| Conferencing | 102 | 16 | 59 |
| Other needs | 8 | 32 | 7 |
| Very few responses to “Other needs” given; one example: training of field workers and students in field identification. |  |  |  |

Survey participants were also asked to give their opinions in a text field on research priorities for vector biology over the next 5-10 years. Answers varied but some key areas were identified: 1) Vector interactions with hosts and pathogens, including vector competence and transmission; 2) Insecticide resistance and novel insecticides; 3) Ecology and behavior, including of infected vectors, introduction of vectors etc.; 4) Vector control, novel control measures and surveillance; 5) Vaccines, including anti-vector vaccines; 6) Modelling; 7) Vector genomics/genetics and bioinformatics. Although no survey can be complete, the data presented here yields a valuable picture of the needs and requirements in disease vector biology, especially of European scientists. We thus expect this study to be relevant to stakeholders such as governments, research councils and organizations but also researchers as priorities for future activities such as those planned by INFRAVEC-2 are determined.

## Acknowledgments

We acknowledge the assistance of Sarah J. Plowman of the UK BBSRC for advice in survey design, and survey participants.

---

**S1 Table**. The INFRAVEC-2 Survey Questionnaire, as sent out to participants. A brief description of the INRFAVEC-2 community is given, and the aims of the questionnaire explained.

**S2 Table.**
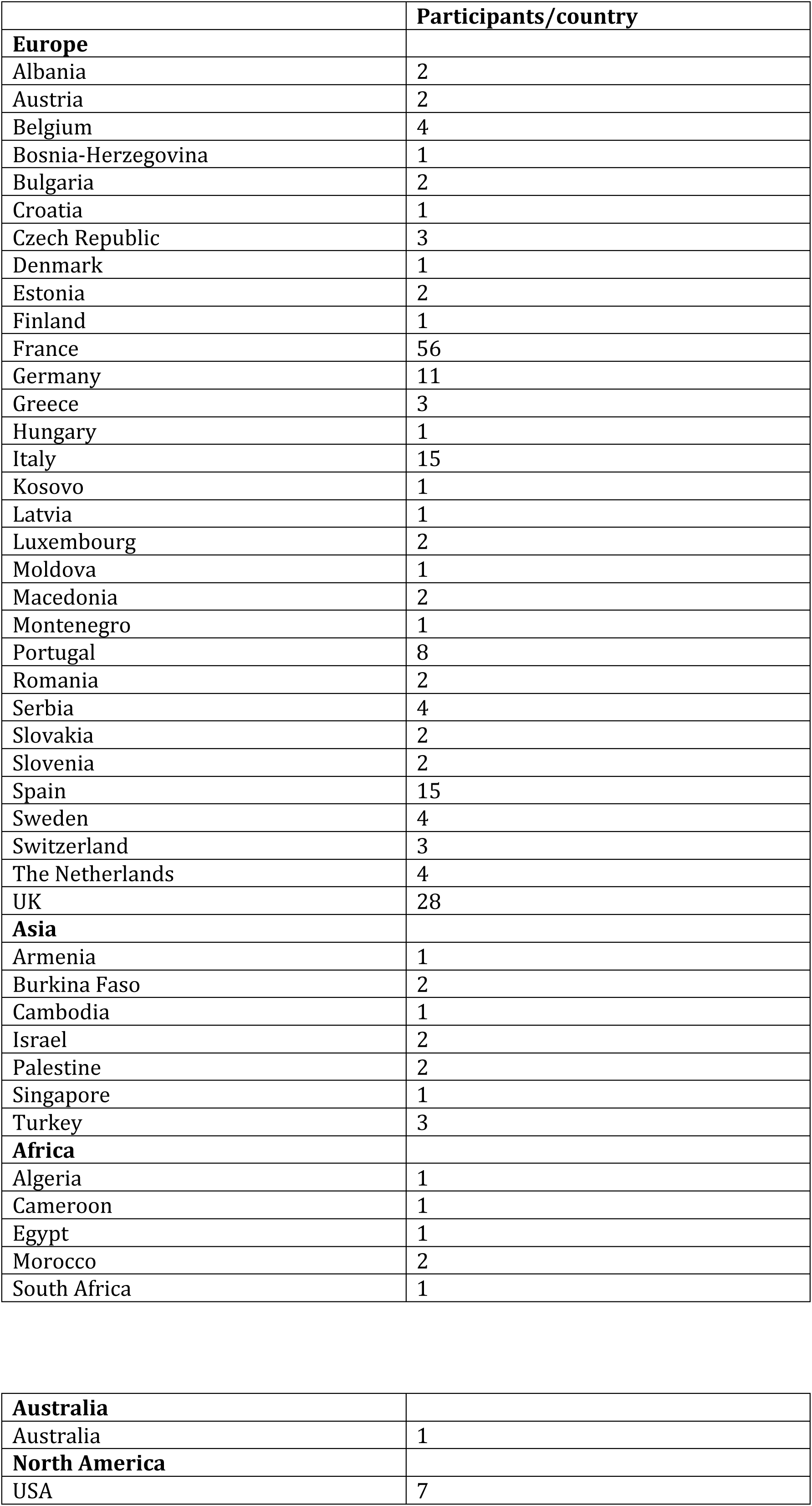
Responses to the INFRAVEC-2 and participation numbers by country, split by continent.

**HORIZON 2020 INFRAVEC-2 Questionnaire**

**Introduction**

A consortium of European institutions based in the former FP7/INFRAVEC project is responding to the new H2020 call “Integrating Activities for Advanced Communities” of the European Research Infrastructures (RI) Programme under the item “Research Infrastructures for the control of vector-borne diseases”. The primary purpose of an “integrated infrastructure” is to provide the EU scientific community with access to its network of RI facilities and services, without charge to the end user, at the state-of-the art premises of participating institutions. Access to the specialized RI enables European researchers and SME to carry out experiments beyond their current capacities.

Building upon the major achievements of FP7/INFRAVEC in forging a European Starting Community of insect vector RI, we worked with EC representatives to generate the current H2020 call for an Advanced Community (AC). With strong commitment obtained from collaborating institutions hosting top-level specialized EU facilities for vector-borne disease (including Institut Pasteur FR, Imperial College UK, Centre de Recerca en Sanitat Animal (IRTA-CReSA) ES, Wageningen University NL, University of Glasgow UK, Institut de Recherche pour le Développement (IRD)-Montpellier FR, Polo d’Innovazione di Genomica, Genetica e Biologia (Polo GGB) IT, Pirbright Institute UK, Max-Planck-Institut für Infektionsbiologie DE, Radboud University Medical Center NL, and EMBL European Bioinformatics Institute DE), the group, chaired by K. Vernick (Institut Pasteur), has been invited to organize the AC.

The RI consortium will provide enabling infrastructures and support for research on disease vectors and their pathogens. We have listed possible RI and services within the following questionnaire, and we would like to solicit as wide as possible feedback from potential users in order to understand the major needs of the vector biology community. Your particular requirements and feedback will have strong impact on how the project will be structured, as this “integrated infrastructure” needs to be tightly tailored to, and inspired by real community needs. Please, take a minute to fill in a short questionnaire (~15 min) that will help us mobilize necessary resources for the future of our community.

Please feel free to forward this email to relevant colleagues. The primary target audience is EU insect vector researchers and SME, but we welcome replies from outside the EU as well. All individual replies and identity information will be kept confidential. Questions can be addressed to

Please complete this form as soon as you can. The survey will close November 25th 2015.

Thank you for your collaboration.

**Q1. Please provide your name and contact details**.

**Title**

_____

**First name**

_____

**Last name**

_____

**Position**

_____

**Organization**

_____

**Country**

_____

**Email (optional)**

Providing an email address is optional, but will permit us to keep you informed about the consortium; contact information will not be shared with other parties.

_____

**Q2. Please identify the arthropod vectors and/or vector borne pathogens that you research**

**Select all that apply**

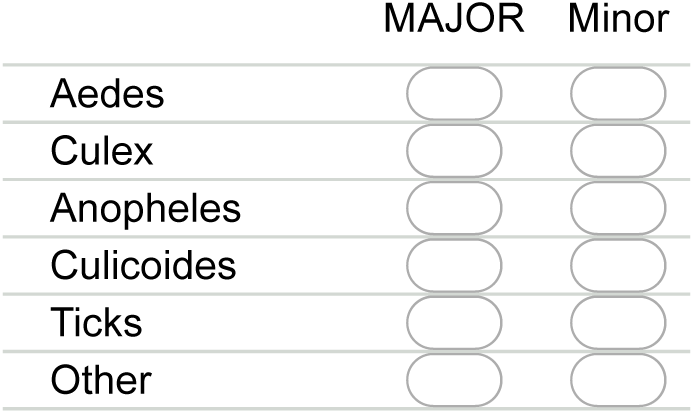

**If other other area**

Please specify below

**Select all that apply**

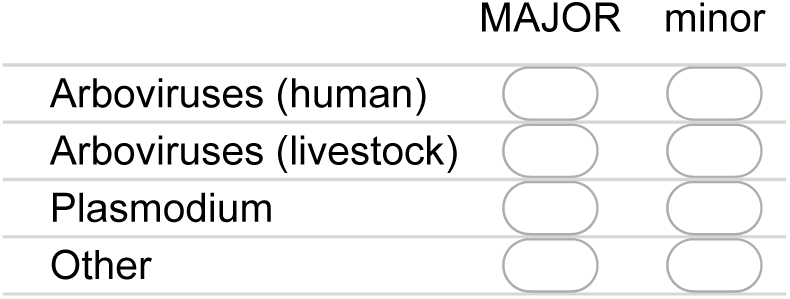

**If other other area**

Please specify below

**Q3. Please select the most relevant areas that describe your research interests from the list below**

**Select all that apply**

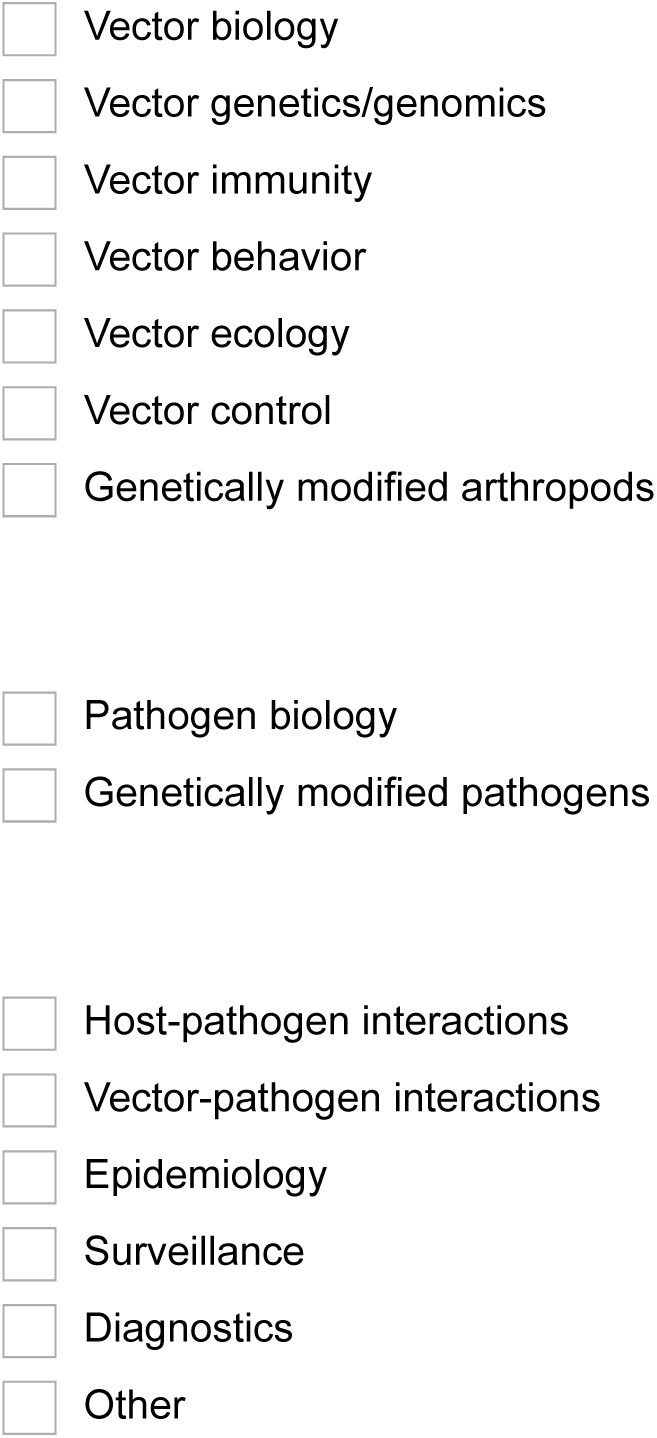

**If other**

Please specify below

**Q4. Does your organization have infrastructure facilities described by the list below?**

**Select all that apply**

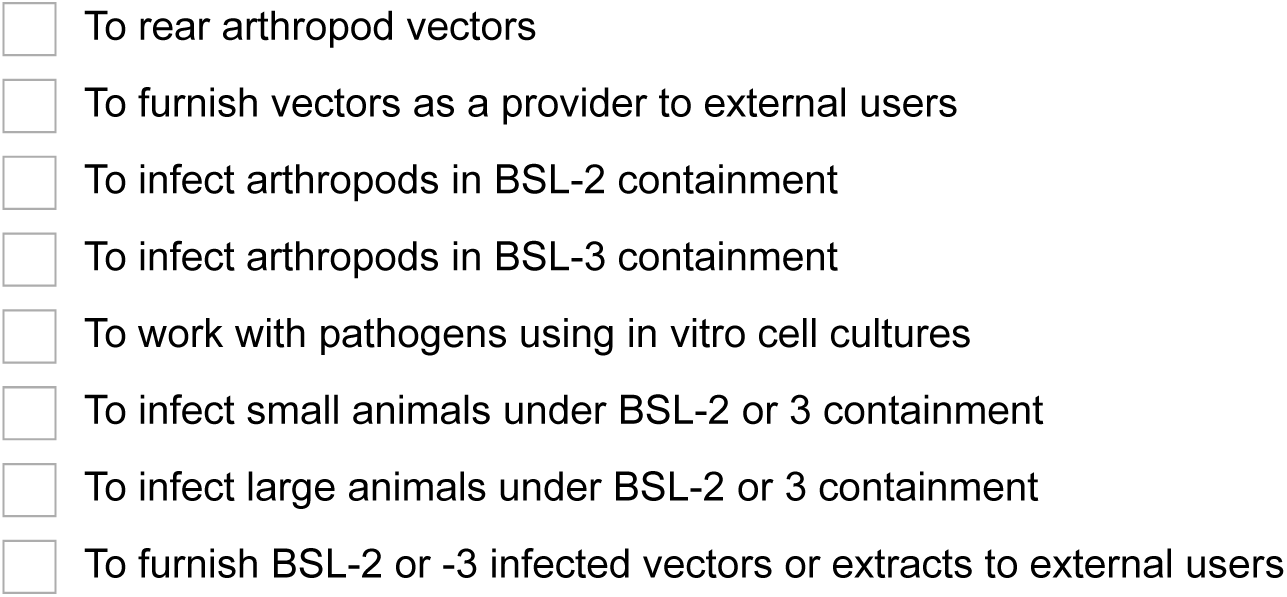

**Q5. Have you ever tried to access BSL-2 or 3 vector research facilities based at organizations other than your own?**

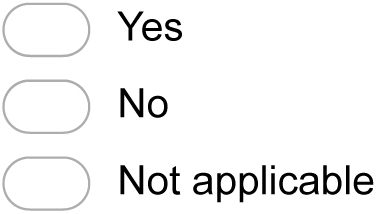

**a. If yes, did the facility have sufficient capacity to accommodate your request in a timely manner?**

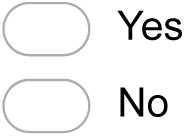

**Q6. Which infrastructure services offered to European users would you be likely to use, with user access costs paid by a Horizon 2020 Research Infrastructure consortium (i.e., at no charge to the end-user). Items provided as user access or custom service, which does not require scientific collaboration with the providing facility**.

**a. VECTOR INFECTION AND VECTOR-PATHOGEN INTERACTIONS. Access to BSL-2 or 3 secure insectary facilities for infection of vectors, or provision of infected vectors or extracts custom-generated by such a facility. Vectors infected by the following pathogens, and for the following research purposes (select all that apply)**.

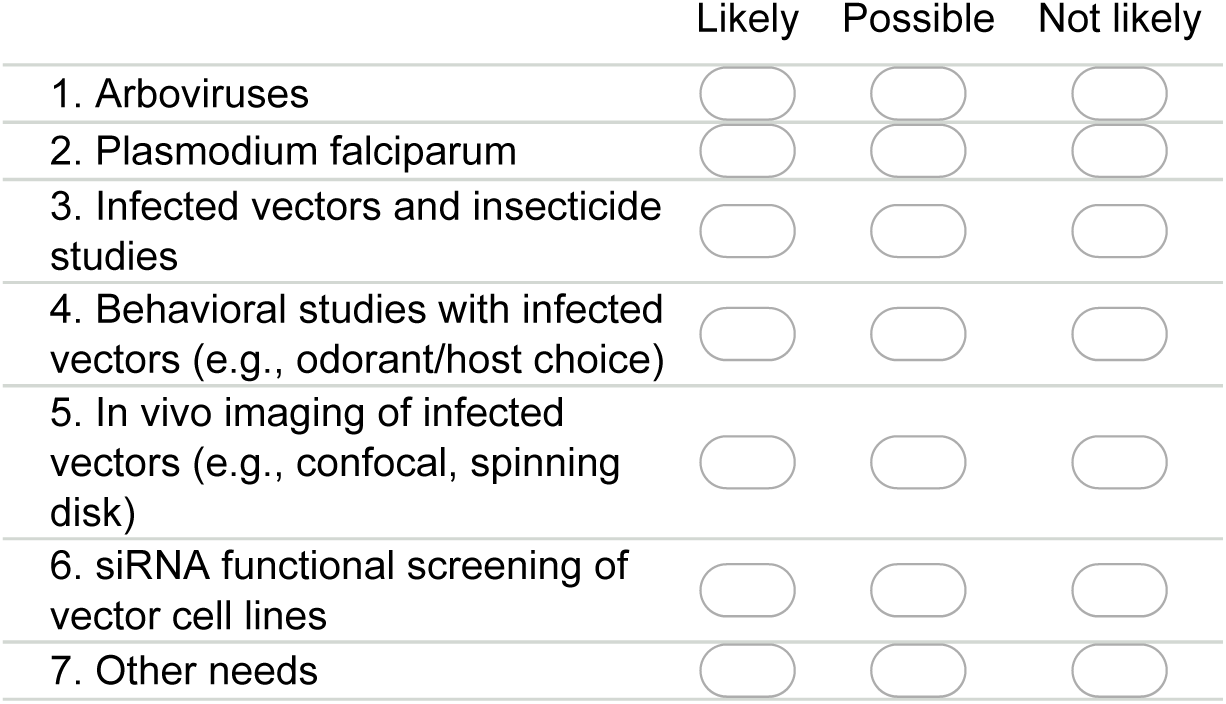

If other needs–please specify below

_____

_____

_____

_____

_____

**Q6. Which infrastructure services offered to European users would you be likely to use, with user access costs paid by a Horizon 2020 Research Infrastructure consortium (i.e., at no charge to the end-user). Items provided as user access or custom service, which does not require scientific collaboration with the providing facility**.

**b. VECTOR GENOMICS AND BIOINFORMATICS. High-throughput genomic services. If desired, with upstream bioinformatic design advice and downstream bioinformatic analysis (select all that apply)**

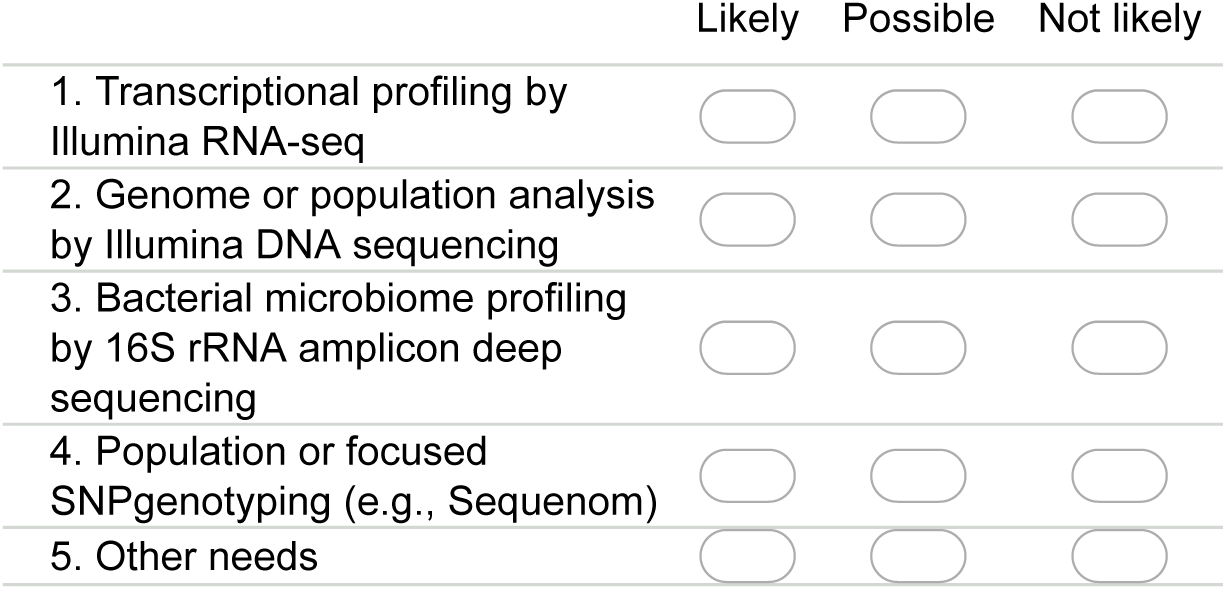

If other needs–please specify below

_____

_____

_____

_____

_____

**Q6. Which infrastructure services offered to European users would you be likely to use, with user access costs paid by a Horizon 2020 Research Infrastructure consortium (i.e., at no charge to the end-user). Items provided as user access or custom service, which does not require scientific collaboration with the providing facility**.

**c. VECTOR GENOME EDITING. Provision of custom genetic modification of your requested target gene or sequence using CRISPR or other technology in vectors (select all that apply). Could also include phenotyping the mutation effect by pathogen challenge under (a) above**.

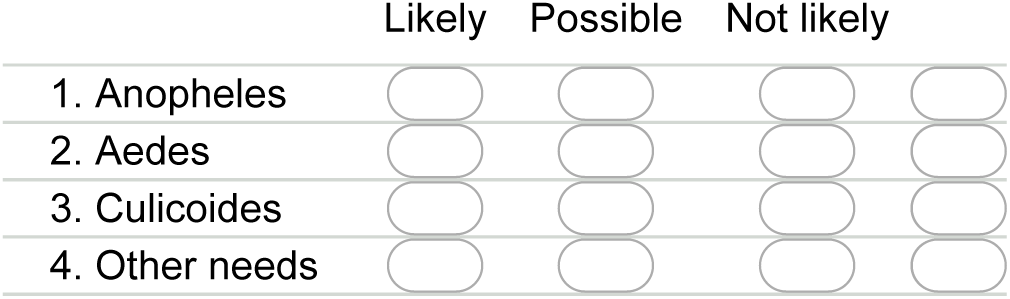

If other needs–please specify below

_____

_____

_____

_____

_____

**Q6. Which infrastructure services offered to European users would you be likely to use, with user access costs paid by a Horizon 2020 Research Infrastructure consortium (i.e., at no charge to the end-user). Items provided as user access or custom service, which does not require scientific collaboration with the providing facility**.

**d. VECTOR ECOLOGY AND BEHAVIOR. Provision of access to facilities or custom-performed assays (select all that apply). Could also include pathogen infection of vectors under (a) above**.

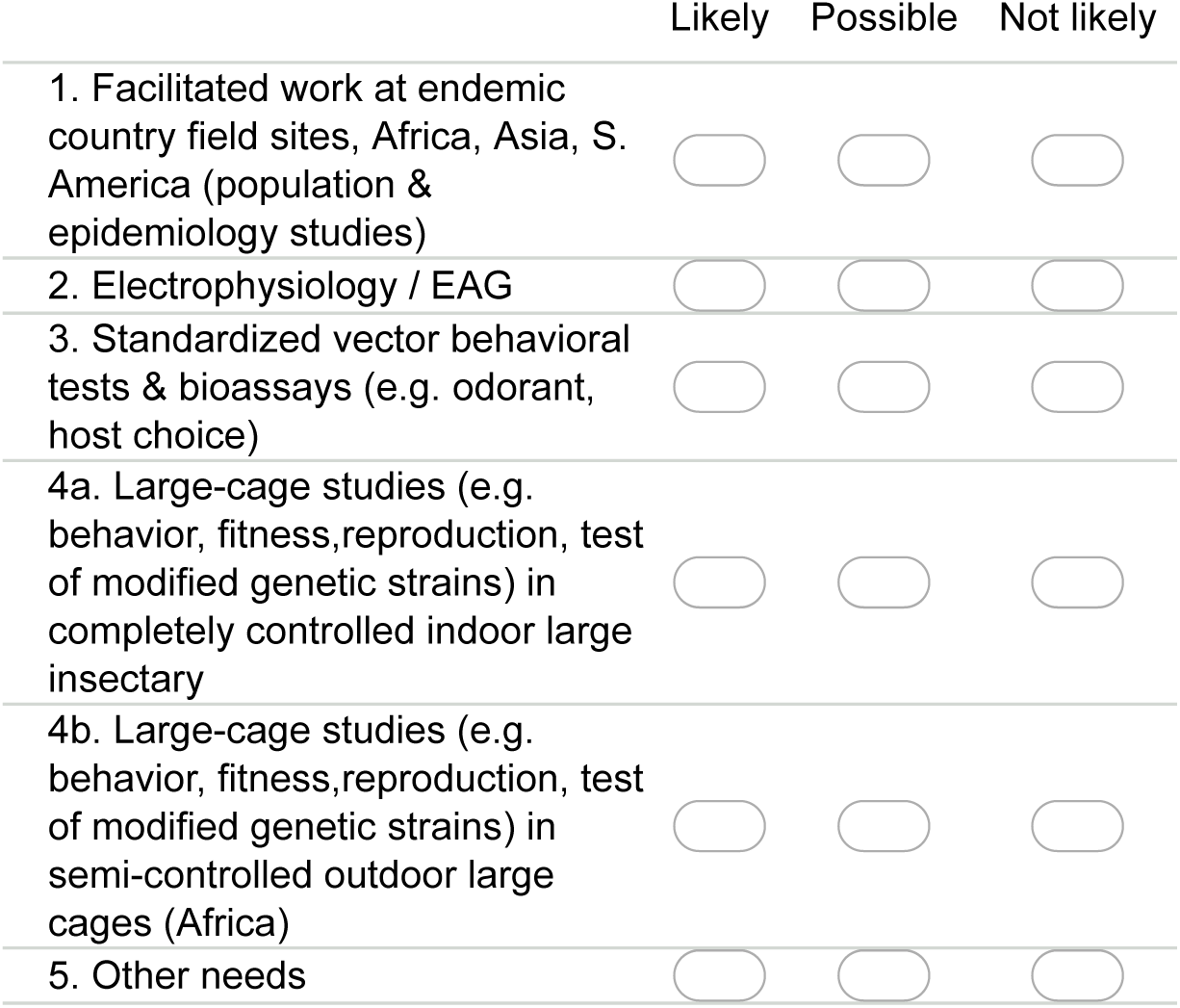

If other needs–please specify below

_____

_____

_____

_____

_____

**Q6. Which infrastructure services offered to European users would you be likely to use, with user access costs paid by a Horizon 2020 Research Infrastructure consortium (i.e., at no charge to the end-user). Items provided as user access or custom service, which does not require scientific collaboration with the providing facility**.

**e. VECTOR BIOLOGY RESOURCES. Provision of vector research resources by request (select all that apply)**.

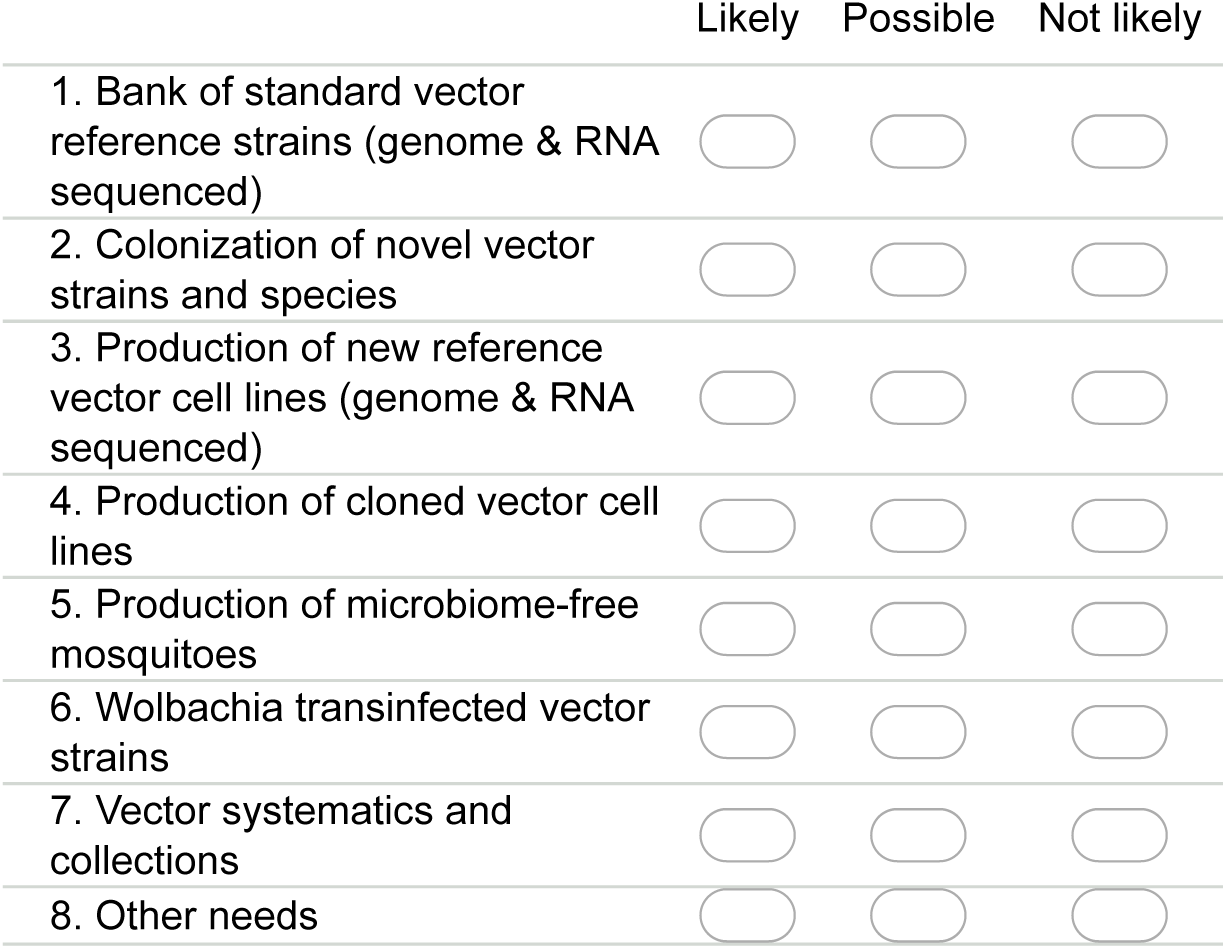

If other needs–please specify below

_____

_____

_____

_____

_____

**Q6. Which infrastructure services offered to European users would you be likely to use, with user access costs paid by a Horizon 2020 Research Infrastructure consortium (i.e., at no charge to the end-user). Items provided as user access or custom service, which does not require scientific collaboration with the providing facility**.

**f. TRAINING AND NETWORKING ACTIVITIES. Promotion of expertise using standardized, comparable practices, scientific exchange**.

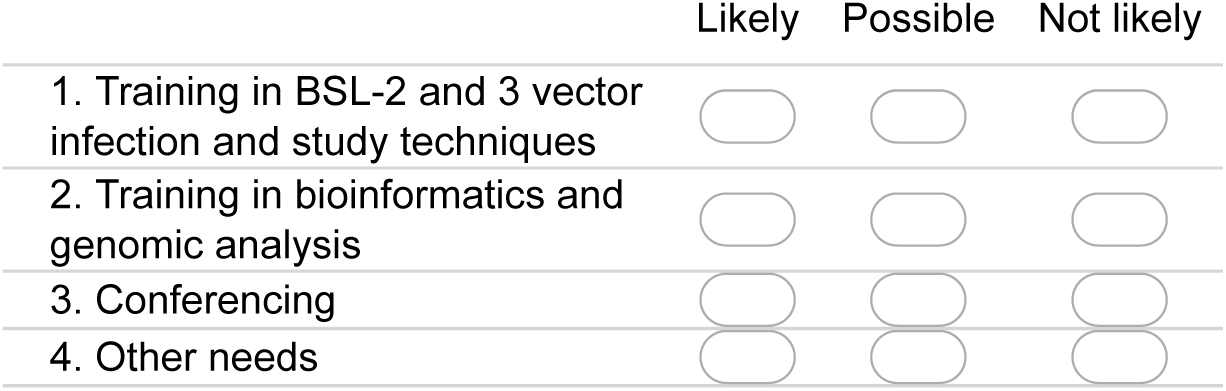

If other needs–please specify below

_____

_____

_____

_____

_____

**Q7. In your opinion, what are the top research priorities (up to 5) in vector biology and/or vector borne disease that need to be addressed in the next 5-10 yrs in the European research context?**

1.

_____

_____

_____

_____

_____

2.

_____

_____

_____

_____

_____

3.

_____

_____

_____

_____

_____

4.

_____

_____

_____

_____

_____

5.

_____

_____

_____

_____

_____

**Thank you for participating!**

**Please use the space provided below to send us any additional feedback on this survey**.

_____

_____

_____

_____

_____

## References

1. Weaver SC, Lecuit M. Chikungunya virus and the global spread of a mosquito-borne disease. N Engl J Med. 2015;372(13):1231–9. doi: 10.1056/NEJMra1406035. PubMed PMID: 25806915.

2. Gatherer D, Kohl A. Zika virus: a previously slow pandemic spreads rapidly through the Americas. J Gen Virol. 2015. doi: 10.1099/jgv.0.000381. PubMed PMID: 26684466.

3. Guzman MG, Harris E. Dengue. Lancet. 2014;385:453–65. doi: 10.1016/S0140-6736(14)60572-9. PubMed PMID: 25230594.

4. Miller LH, Ackerman HC, Su XZ, Wellems TE. Malaria biology and disease pathogenesis: insights for new treatments. Nat Med. 2013;19(2):156–67. doi: 10.1038/nm.3073. PubMed PMID: 23389616.

5. Halbroth BR, Draper SJ. Recent developments in malaria vaccinology. Adv Parasitol. 2015;88:1–49. doi: 10.1016/bs.apar.2015.03.001. PubMed PMID: 25911364.

6. Hoffman SL, Vekemans J, Richie TL, Duffy PE. The march toward malaria vaccines. Vaccine. 2015;33 Suppl 4:D13–23. doi: 10.1016/j.vaccine.2015.07.091. PubMed PMID: 26324116.

7. Thomas SJ, Rothman AL. Trials and tribulations on the path to developing a dengue vaccine. Vaccine. 2015;33 Suppl 4:D24–31. doi: 10.1016/j.vaccine.2015.05.095. PubMed PMID: 26122583.

8. Schwartz LM, Halloran ME, Durbin AP, Longini IM, Jr.. The dengue vaccine pipeline: Implications for the future of dengue control. Vaccine. 2015;33(29):3293–8. doi: 10.1016/j.vaccine.2015.05.010. PubMed PMID: 25989449; PubMed Central PMCID: PMCPMC4470297.

9. Neafsey DE, Juraska M, Bedford T, Benkeser D, Valim C, Griggs A, et al. Genetic Diversity and Protective Efficacy of the RTS,S/AS01 Malaria Vaccine. N Engl J Med. 2015;373(21):2025–37. doi: 10.1056/NEJMoa1505819. PubMed PMID: 26488565.

10. Newby G, Hwang J, Koita K, Chen I, Greenwood B, von Seidlein L, et al. Review of mass drug administration for malaria and its operational challenges. Am J Trop Med Hyg. 2015;93(1):125–34. doi: 10.4269/ajtmh.14-0254. PubMed PMID: 26013371; PubMed Central PMCID: PMCPMC4497884.

11. Wells TN, Hooft van Huijsduijnen R, Van Voorhis WC. Malaria medicines: a glass half full? Nat Rev Drug Discov. 2015;14(6):424–42. doi: 10.1038/nrd4573. PubMed PMID: 26000721.

12. Sinha S, Medhi B, Sehgal R. Challenges of drug-resistant malaria. Parasite. 2014;21:61. doi: 10.1051/parasite/2014059. PubMed PMID: 25402734; PubMed Central PMCID: PMCPMC4234044.

13. Lim SP, Noble CG, Shi PY. The dengue virus NS5 protein as a target for drug discovery. Antiviral Res. 2015;119:57–67. doi: 10.1016/j.antiviral.2015.04.010. PubMed PMID: 25912817.

14. Chen YL, Yokokawa F, Shi PY. The search for nucleoside/nucleotide analog inhibitors of dengue virus. Antiviral Res. 2015;122:12–9. doi: 10.1016/j.antiviral.2015.07.010. PubMed PMID: 26241002.

15. Lim SP, Wang QY, Noble CG, Chen YL, Dong H, Zou B, et al. Ten years of dengue drug discovery: progress and prospects. Antiviral Res. 2013;100(2):500–19. doi: 10.1016/j.antiviral.2013.09.013. PubMed PMID: 24076358.

16. Xie X, Zou J, Wang QY, Shi PY. Targeting dengue virus NS4B protein for drug discovery. Antiviral Res. 2015;118:39–45. doi: 10.1016/j.antiviral.2015.03.007. PubMed PMID: 25796970.

17. Ahola T, Courderc T, Ng LF, Hallengard D, Powers A, Lecuit M, et al. Therapeutics and vaccines against chikungunya virus. Vector Borne Zoonotic Dis. 2015;15(4):250–7. doi: 10.1089/vbz.2014.1681. PubMed PMID: 25897811.

18. Kortekaas J. One Health approach to Rift Valley fever vaccine development. Antiviral Res. 2014;106:24–32. doi: 10.1016/j.antiviral.2014.03.008. PubMed PMID: 24681125.

19. Mansfield KL, Banyard AC, McElhinney L, Johnson N, Horton DL, Hernandez-Triana LM, et al. Rift Valley fever virus: A review of diagnosis and vaccination, and implications for emergence in Europe. Vaccine. 2015;33(42):5520–31. doi: 10.1016/j.vaccine.2015.08.020. PubMed PMID: 26296499.

20. Ocampo CB, Mina NJ, Carabali M, Alexander N, Osorio L. Reduction in dengue cases observed during mass control of Aedes (Stegomyia) in street catch basins in an endemic urban area in Colombia. Acta Trop. 2014;132:15–22. doi: 10.1016/j.actatropica.2013.12.019. PubMed PMID: 24388794; PubMed Central PMCID: PMCPMC4654410.

21. Abad-Franch F, Zamora-Perea E, Ferraz G, Padilla-Torres SD, Luz SL. Mosquito-disseminated pyriproxyfen yields high breeding-site coverage and boosts juvenile mosquito mortality at the neighborhood scale. PLoS Negl Trop Dis. 2015;9(4):e0003702. doi: 10.1371/journal.pntd.0003702. PubMed PMID: 25849040; PubMed Central PMCID: PMCPMC4388722.

22. Harris C, Kihonda J, Lwetoijera D, Dongus S, Devine G, Majambere S. A simple and efficient tool for trapping gravid Anopheles at breeding sites. Parasit Vectors. 2011;4:125. doi: 10.1186/1756-3305-4-125. PubMed PMID: 21722391; PubMed Central PMCID: PMCPMC3141746.

23. Salem OA, Khadijetou ML, Moina MH, Lassana K, Sebastien B, Ousmane F, et al. Characterization of anopheline (Diptera: Culicidae) larval habitats in Nouakchott, Mauritania. J Vector Borne Dis. 2013;50(4):302–6. PubMed PMID: 24499854.

24. Helinski ME, Nuwa A, Protopopoff N, Feldman M, Ojuka P, Oguttu DW, et al. Entomological surveillance following a long-lasting insecticidal net universal coverage campaign in Midwestern Uganda. Parasit Vectors. 2015;8:458. doi: 10.1186/s13071-015-1060-6. PubMed PMID: 26382583; PubMed Central PMCID: PMCPMC4574096.

25. Kawada H, Dida GO, Ohashi K, Kawashima E, Sonye G, Njenga SM, et al. A small-scale field trial of pyriproxyfen-impregnated bed nets against pyrethroid-resistant Anopheles gambiae s.s. in western Kenya. PLoS One. 2014;9(10):e111195. doi: 10.1371/journal.pone.0111195. PubMed PMID: 25333785; PubMed Central PMCID: PMCPMC4205095.

26. Sokhna C, Ndiath MO, Rogier C. The changes in mosquito vector behaviour and the emerging resistance to insecticides will challenge the decline of malaria. Clin Microbiol Infect. 2013;19(10):902–7. doi: 10.1111/1469-0691.12314. PubMed PMID: 23910459.

27. Gurtler RE, Garelli FM, Coto HD. Effects of a five-year citywide intervention program to control Aedes aegypti and prevent dengue outbreaks in northern Argentina. PLoS Negl Trop Dis. 2009;3(4):e427. doi: 10.1371/journal.pntd.0000427. PubMed PMID: 19399168; PubMed Central PMCID: PMCPMC2669131.

28. Padilla-Torres SD, Ferraz G, Luz SL, Zamora-Perea E, Abad-Franch F. Modeling dengue vector dynamics under imperfect detection: three years of site-occupancy by Aedes aegypti and Aedes albopictus in urban Amazonia. PLoS One. 2013;8(3):e58420. doi: 10.1371/journal.pone.0058420. PubMed PMID: 23472194; PubMed Central PMCID: PMCPMC3589427.

29. Norris LC, Main BJ, Lee Y, Collier TC, Fofana A, Cornel AJ, et al. Adaptive introgression in an African malaria mosquito coincident with the increased usage of insecticide-treated bed nets. Proc Natl Acad Sci U S A. 2015;112(3):815–20. doi: 10.1073/pnas.1418892112. PubMed PMID: 25561525; PubMed Central PMCID: PMCPMC4311837.

30. Yewhalaw D, Asale A, Tushune K, Getachew Y, Duchateau L, Speybroeck N. Bio-efficacy of selected long-lasting insecticidal nets against pyrethroid resistant Anopheles arabiensis from South-Western Ethiopia. Parasit Vectors. 2012;5:159. doi: 10.1186/1756-3305-5-159. PubMed PMID: 22871143; PubMed Central PMCID: PMCPMC3485103.

31. Ngufor C, N’Guessan R, Fagbohoun J, Subramaniam K, Odjo A, Fongnikin A, et al. Insecticide resistance profile of Anopheles gambiae from a phase II field station in Cove, southern Benin: implications for the evaluation of novel vector control products. Malar J. 2015;14(1):464. doi: 10.1186/s12936-015-0981-z. PubMed PMID: 26581678; PubMed Central PMCID: PMCPMC4652434.

32. Sande S, Zimba M, Chinwada P, Masendu HT, Mazando S, Makuwaza A. The emergence of insecticide resistance in the major malaria vector Anopheles funestus (Diptera: Culicidae) from sentinel sites in Mutare and Mutasa Districts, Zimbabwe. Malar J. 2015;14(1):466. doi: 10.1186/s12936-015-0993-8. PubMed PMID: 26589891; PubMed Central PMCID: PMCPMC4654866.

33. Djogbenou LS, Assogba B, Essandoh J, Constant EA, Makoutode M, Akogbeto M, et al. Estimation of allele-specific Ace-1 duplication in insecticide-resistant Anopheles mosquitoes from West Africa. Malar J. 2015;14(1):507. doi: 10.1186/s12936-015-1026-3. PubMed PMID: 26682913; PubMed Central PMCID: PMCPMC4683970.

34. Alout H, Labbe P, Berthomieu A, Makoundou P, Fort P, Pasteur N, et al. High chlorpyrifos resistance in Culex pipiens mosquitoes: strong synergy between resistance genes. Heredity (Edinb). 2015. doi: 10.1038/hdy.2015.92. PubMed PMID: 26463842.

35. Misra BR, Gore M. Malathion Resistance Status and Mutations in Acetylcholinesterase Gene (Ace) in Japanese Encephalitis and Filariasis Vectors from Endemic Area in India. J Med Entomol. 2015;52(3):442–6. doi: 10.1093/jme/tjv015. PubMed PMID: 26334819.

36. Giraldo-Calderon GI, Emrich SJ, MacCallum RM, Maslen G, Dialynas E, Topalis P, et al. VectorBase: an updated bioinformatics resource for invertebrate vectors and other organisms related with human diseases. Nucleic Acids Res. 2015;43(Database issue):D707–13. doi: 10.1093/nar/gku1117. PubMed PMID: 25510499; PubMed Central PMCID: PMCPMC4383932.

37. Kean J, Rainey SM, McFarlane M, Donald CL, Schnettler E, Kohl A, et al. Fighting Arbovirus Transmission: Natural and Engineered Control of Vector Competence in Aedes Mosquitoes. Insects. 2015;6(1):236–78. doi: 10.3390/insects6010236. PubMed PMID: 26463078; PubMed Central PMCID: PMCPMC4553541.

38. Fraser MJ, Jr. Insect transgenesis: current applications and future prospects. Annu Rev Entomol. 2012;57:267–89. doi: 10.1146/annurev.ento.54.110807.090545. PubMed PMID: 22149266.

39. Nolan T, Papathanos P, Windbichler N, Magnusson K, Benton J, Catteruccia F, et al. Developing transgenic Anopheles mosquitoes for the sterile insect technique. Genetica. 2011;139(1):33–9. doi: 10.1007/s10709-010-9482-8. PubMed PMID: 20821345.

40. Alphey L. Genetic control of mosquitoes. Annu Rev Entomol. 2014;59:205–24. Epub 2013/10/29. doi: 10.1146/annurev-ento-011613-162002. PubMed PMID: 24160434.

41. Alphey N, Bonsall MB. Interplay of population genetics and dynamics in the genetic control of mosquitoes. J R Soc Interface. 2014;11(93):20131071. Epub 2014/02/14. doi: 10.1098/rsif.2013.1071 rsif.2013.1071 [pii]. PubMed PMID: 24522781; PubMed Central PMCID: PMC3928937.

42. Franz AW, Clem RJ, Passarelli AL. Novel Genetic and Molecular Tools for the Investigation and Control of Dengue Virus Transmission by Mosquitoes. Curr Trop Med Rep. 2014;1(1):21–31. doi: 10.1007/s40475-013-0007-2. PubMed PMID: 24693489; PubMed Central PMCID: PMCPMC3969738.

43. Alphey L, McKemey A, Nimmo D, Neira Oviedo M, Lacroix R, Matzen K, et al. Genetic control of Aedes mosquitoes. Pathog Glob Health. 2013;107(4):170–9. Epub 2013/07/03. doi: 10.1179/2047773213Y.0000000095. PubMed PMID: 23816508.

44. Hegde S, Rasgon JL, Hughes GL. The microbiome modulates arbovirus transmission in mosquitoes. Curr Opin Virol. 2015;15:97–102. doi: 10.1016/j.coviro.2015.08.011. PubMed PMID: 26363996.

45. Clayton AM, Dong Y, Dimopoulos G. The Anopheles innate immune system in the defense against malaria infection. Journal of innate immunity. 2014;6(2):169–81. doi: 10.1159/000353602. PubMed PMID: 23988482; PubMed Central PMCID: PMCPMC3939431.

46. Jupatanakul N, Sim S, Dimopoulos G. The insect microbiome modulates vector competence for arboviruses. Viruses. 2014;6(11):4294–313. doi: 10.3390/v6114294. PubMed PMID: 25393895; PubMed Central PMCID: PMCPMC4246223.

47. Bolling BG, Olea-Popelka FJ, Eisen L, Moore CG, Blair CD. Transmission dynamics of an insect-specific flavivirus in a naturally infected Culex pipiens laboratory colony and effects of co-infection on vector competence for West Nile virus. Virology. 2012;427(2):90–7. doi: 10.1016/j.virol.2012.02.016. PubMed PMID: 22425062; PubMed Central PMCID: PMCPMC3329802.

48. Mosimann AL, Bordignon J, Mazzarotto GC, Motta MC, Hoffmann F, Santos CN. Genetic and biological characterization of a densovirus isolate that affects dengue virus infection. Mem Inst Oswaldo Cruz. 2011;106(3):285–92. PubMed PMID: 21655815.

49. Rainey SM, Shah P, Kohl A, Dietrich I. Understanding the Wolbachia-mediated inhibition of arboviruses in mosquitoes: progress and challenges. J Gen Virol. 2014;95(Pt 3):517–30. Epub 2013/12/18. doi: 10.1099/vir.0.057422-0 vir.0.057422-0 [pii]. PubMed PMID: 24343914.

50. Iturbe-Ormaetxe I, Walker T, Sl ON. Wolbachia and the biological control of mosquito-borne disease. EMBO Rep. 2011;12(6):508–18. Epub 2011/05/07. doi: embor201184 [pii]10.1038/embor.2011.84. PubMed PMID: 21546911.

51. Johnson KN. The Impact of Wolbachia on Virus Infection in Mosquitoes. Viruses. 2015;7(11):5705–17. doi: 10.3390/v7112903. PubMed PMID: 26556361; PubMed Central PMCID: PMCPMC4664976.

52. Lambrechts L, Ferguson NM, Harris E, Holmes EC, McGraw EA, O’Neill SL, et al. Assessing the epidemiological effect of wolbachia for dengue control. Lancet Infect Dis. 2015;15(7):862–6. doi: 10.1016/S1473-3099(15)00091-2. PubMed PMID: 26051887.

53. http://ec.europa.eu/research/infrastructures/index_en.cfm.

54. Angelini R, Finarelli AC, Angelini P, Po C, Petropulacos K, Macini P, et al. An outbreak of chikungunya fever in the province of Ravenna, Italy. Euro Surveill. 2007;12(9):E070906 1. Epub 2007/09/29. doi: 2260 [pii]. PubMed PMID: 17900424.

55. Burt FJ, Rolph MS, Rulli NE, Mahalingam S, Heise MT. Chikungunya: a re-emerging virus. Lancet. 2012;379(9816):662–71. Epub 2011/11/22. doi: S0140-6736(11)60281-X [pii] 10.1016/S0140-6736(11)60281-X. PubMed PMID: 22100854.

56. Coffey LL, Failloux AB, Weaver SC. Chikungunya virus-vector interactions. Viruses. 2014;6(11):4628–63. doi: 10.3390/v6114628. PubMed PMID: 25421891; PubMed Central PMCID: PMC4246241.

57. Weaver SC, Forrester NL. Chikungunya: Evolutionary history and recent epidemic spread. Antiviral Res. 2015;120:32–9. doi: 10.1016/j.antiviral.2015.04.016. PubMed PMID: 25979669.

58. Lambrechts L, Scott TW, Gubler DJ. Consequences of the expanding global distribution of Aedes albopictus for dengue virus transmission. PLoS Negl Trop Dis. 2010;4(5):e646. doi: 10.1371/journal.pntd.0000646. PubMed PMID: 20520794; PubMed Central PMCID: PMC2876112.

59. Paupy C, Delatte H, Bagny L, Corbel V, Fontenille D. Aedes albopictus, an arbovirus vector: From the darkness to the light. Microbes Infect. 2009. Epub 2009/05/20. doi: S1286-4579(09)00105-1 [pii] 10.1016/j.micinf.2009.05.005. PubMed PMID: 19450706.

60. Kraemer MU, Sinka ME, Duda KA, Mylne AQ, Shearer FM, Barker CM, et al. The global distribution of the arbovirus vectors Aedes aegypti and Ae. albopictus. Elife. 2015;4:e08347. doi: 10.7554/eLife.08347. PubMed PMID: 26126267; PubMed Central PMCID: PMCPMC4493616.

61. Beer M, Conraths FJ, van der Poel WH. ‘Schmallenberg virus’–a novel orthobunyavirus emerging in Europe. Epidemiol Infect. 2013;141(1):1–8. Epub 2012/10/11. doi: 10.1017/S0950268812002245S0950268812002245 [pii]. PubMed PMID: 23046921.

62. Powers AM. Risks to the Americas Associated with the Continued Expansion of Chikungunya Virus. J Gen Virol. 2014. doi: 10.1099/vir.0.070136-0. PubMed PMID: 25239764.

63. Carpenter S, Wilson A, Mellor PS. Culicoides and the emergence of bluetongue virus in northern Europe. Trends Microbiol. 2009;17(4):172–8. Epub 2009/03/21. doi: S0966-842X(09)00040-7 [pii] 10.1016/j.tim.2009.01.001. PubMed PMID: 19299131.

64. Schotthoefer AM, Frost HM. Ecology and Epidemiology of Lyme Borreliosis. Clinics in laboratory medicine. 2015;35(4):723–43. doi: 10.1016/j.cll.2015.08.003. PubMed PMID: 26593254.

65. Ergonul O. Crimean-Congo hemorrhagic fever virus: new outbreaks, new discoveries. Curr Opin Virol. 2012;2(2):215–20. Epub 2012/04/10. doi: 10.1016/j.coviro.2012.03.001S1879-6257(12)00044-2 [pii]. PubMed PMID: 22482717.

66. Papa A, Mirazimi A, Koksal I, Estrada-Pena A, Feldmann H. Recent advances in research on Crimean-Congo hemorrhagic fever. J Clin Virol. 2015;64:137–43. doi: 10.1016/j.jcv.2014.08.029. PubMed PMID: 25453328; PubMed Central PMCID: PMCPMC4346445.

67. Cirimotich CM, Ramirez JL, Dimopoulos G. Native microbiota shape insect vector competence for human pathogens. Cell Host Microbe. 2011;10(4):307–10. doi: 10.1016/j.chom.2011.09.006. PubMed PMID: 22018231; PubMed Central PMCID: PMCPMC3462649.

68. Arensburger P, Megy K, Waterhouse RM, Abrudan J, Amedeo P, Antelo B, et al. Sequencing of Culex quinquefasciatus establishes a platform for mosquito comparative genomics. Science. 2010;330(6000):86–8. Epub 2010/10/12. doi: 330/6000/86 [pii] 10.1126/science.1191864. PubMed PMID: 20929810.

69. Nene V, Wortman JR, Lawson D, Haas B, Kodira C, Tu ZJ, et al. Genome sequence of Aedes aegypti, a major arbovirus vector. Science. 2007;316(5832):1718–23. PubMed PMID: 17510324.

70. Holt RA, Subramanian GM, Halpern A, Sutton GG, Charlab R, Nusskern DR, et al. The genome sequence of the malaria mosquito Anopheles gambiae. Science. 2002;298(5591):129–49. PubMed PMID: 12364791.

71. Dong Y, Das S, Cirimotich C, Souza-Neto JA, McLean KJ, Dimopoulos G. Engineered anopheles immunity to Plasmodium infection. PLoS Pathog. 2011;7(12):e1002458. Epub 2012/01/05. doi: 10.1371/journal.ppat.1002458 PPATHOGENS-D-11-01314 [pii]. PubMed PMID: 22216006; PubMed Central PMCID: PMC3245315.

72. Isaacs AT, Li F, Jasinskiene N, Chen X, Nirmala X, Marinotti O, et al. Engineered resistance to Plasmodium falciparum development in transgenic Anopheles stephensi. PLoS Pathog. 2011;7(4):e1002017. Epub 2011/05/03. doi: 10.1371/journal.ppat.1002017. PubMed PMID: 21533066; PubMed Central PMCID: PMC3080844.

73. Carvalho DO, McKemey AR, Garziera L, Lacroix R, Donnelly CA, Alphey L, et al. Suppression of a Field Population of Aedes aegypti in Brazil by Sustained Release of Transgenic Male Mosquitoes. PLoS Negl Trop Dis. 2015;9(7):e0003864. doi: 10.1371/journal.pntd.0003864. PubMed PMID: 26135160; PubMed Central PMCID: PMCPMC4489809.

74. Hammond A, Galizi R, Kyrou K, Simoni A, Siniscalchi C, Katsanos D, et al. A CRISPR-Cas9 gene drive system targeting female reproduction in the malaria mosquito vector Anopheles gambiae. Nat Biotechnol. 2016;34(1):78–83. doi: 10.1038/nbt.3439. PubMed PMID: 26641531.

75. Gantz VM, Jasinskiene N, Tatarenkova O, Fazekas A, Macias VM, Bier E, et al. Highly efficient Cas9-mediated gene drive for population modification of the malaria vector mosquito Anopheles stephensi. Proc Natl Acad Sci U S A. 2015;112(49):E6736–43. doi: 10.1073/pnas.1521077112. PubMed PMID: 26598698; PubMed Central PMCID: PMCPMC4679060.

